# Effect of Pressure on the Conformational Landscape of Human *γ*D-crystallin from Replica Exchange Molecular Dynamics Simulations

**DOI:** 10.1101/2024.01.07.574493

**Authors:** Arlind Kacirani, Betül Uralcan, Tiago S. Domingues, Amir Haji-Akbari

## Abstract

Human *γ*D-crystallin belongs to a crucial family of proteins known as crystallins located in fiber cells of the human lens. Since crystallins do not undergo any turnover after birth, they need to possess remarkable thermodynamic stability. However, their sporadic misfolding and aggregation, triggered by environmental perturbations or genetic mutations, constitute the molecular basis of cataracts, which is the primary cause of blindness in the globe according to the World Health Organization. Here, we investigate the impact of high pressure on the conformational landscape of the wild-type H*γ*D-crystallin using replica exchange molecular dynamics simulations augmented with principal component analysis. We find pressure to have a modest impact on global measures of protein stability, such as root mean square displacement and radius of gyration. Upon projecting our trajectories along the first two principal components from Pca, however, we observe the emergence of distinct free energy basins at high pressures. By screening local order parameters previously shown or hypothesized as markers of H*γ*D-crystallin stability, we establish correlations between a tyrosine-tyrosine aromatic contact within the N-terminal domain and the protein’s end-to-end distance with projections along the first and second principal components, respectively. Furthermore, we observe the simultaneous contraction of the hydrophobic core and its intrusion by water molecules. This exploration sheds light on the intricate responses of H*γ*D-crystallin to elevated pressures, offering insights into potential mechanisms underlying its stability and susceptibility to environmental perturbations, crucial for understanding cataract formation.

## I. INTRODUCTION

Cataract is the primary cause of blindness worldwide and is the leading factor contributing to vision impairment in the United States.^1^ This condition is characterized by the accumulation of heterogeneous and opaque aggregates formed by a family of mammalian lens proteins referred to as crystallins,^2,3^ which are categorized into three groups based on molecular weight and function. *α*crystallins, which possess higher molecular weights, are heat shock proteins that function as chaperones guiding the folding of *β*and *γ*-crystallins that are responsible for maintaining lens transparency and a high refractive index. Due to lack of turnover post-birth, crystallins require high thermodynamic stability to remain folded, and form stable aqueous solutions at concentrations surpassing 400 g/L.^4^ Specifically, human *γ*Dcrystallin (H*γ*D-crys), a monomeric lens protein comprised of 173 residues, represents a significant component of the human lens whose aggregation contributes to agerelated cataracts.^5,6^ Its secondary structure mainly consists of *β*-sheets organized into two homologous domains (Figure 1A). Each domain contains two Greek-key motifs forming a *β*-sandwich of eight intercalated *β*-strands.^7^

**FIG. 1.**
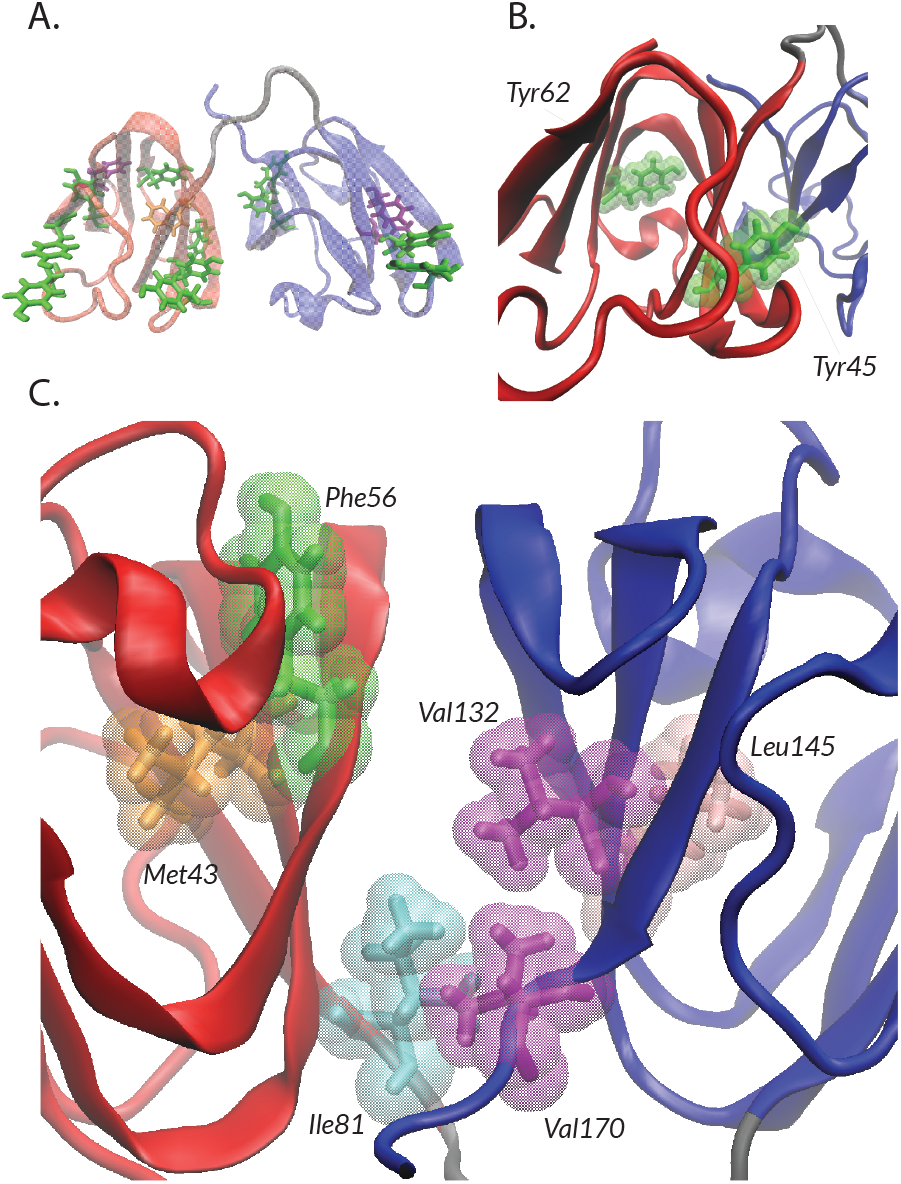
(A) A schematic illustration of H*γ*D-crys with N- and C-terminals depicted in red and blue, respectively, also highlighting the network of relevant aromatic residues. (B) A closer view of the N-terminal domain depicting the van der Waals volumes of tyrosines at positions 45 and 62. (C) Zoomed-in rendering of H*γ*D-crys interdomain interface highlighting the six hydrophobic residues: Met43, Phe56 and Ile81 in N-td, and Val132, Leu145 and Val170 in C-td.

While crystallin aggregation represents a crucial aspect of cataract formation, significant gaps remain in our understanding of the precise molecular mechanisms underlying this phenomenon. When it comes to H*γ*D-crys, an aggregation mechanism that involves domain swapping at the hydrophobic interdomain interface followed by partial unfolding of the N-terminal domain (N-td) has been proposed,^5,8,9^ a proposition supported by single molecule atomic force microscopy (AFM)^10^ and nuclear magnetic resonance (NMR)^11^ experiments, as well as atomistic classical molecular dynamics (Md) simulations.^6^ The structural changes resulting from physicochemical environmental perturbations– such as acidic pH, ultraviolet light, oxidative damage, heating or freezing–^2,6,9,12^ as well as specific amino-acid substitutions,^7,11,13,14^ reportedly contribute to favoring the formation of such aggregation-prone conformations. Therefore, the identification and comprehension of potential initial denaturation sites as early indicators of aggregation are pivotal for enhancing our molecular-level understanding of the mechanism of cataract formation.

Current evidence points to the importance of the aromatic side chain contacts within the N-td (Fig. 1A) and the hydrophobic nature of the interdomain interface (Fig. 1C) to be key markers in the initial denaturation of H*γ*D-crys.^5,15–17^ For instance, alanine substitutions of aromatic residues in H*γ*D-crys N-td, when expressed in *E. coli*, exhibited reduced thermal stability and faster unfolding compared to the wild-type protein.^17^ This underscores the pivotal role of N-td aromatic contacts in maintaining the protein’s structural integrity. In fact, the importance of aromatic pair interactions in folding and stability of a diverse range of proteins is supported by a strong body of experimental evidence,^18–20^ especially in the case of proteins rich in *β*-sheet content.^21–23^ Figure 1A illustrates the structure of H*γ*D-crys, showcasing its N- and C-terminal domains in red and blue, respectively. This visualization highlights the residue pairs engaged in aromatic-aromatic interactions, a critical aspect known to uphold the protein’s folding stability, as previously reported. More specifically, tyrosines emerge as significant contributors to protein structural stability compared to other residues,^24^ due to their ability to form aromatic *π*− *π* contacts, strong hydrogen bonds through the phenol group and other interactions through their multiple side-chain binding modes, as confirmed both experimentally and computationally.^25,26^

Additionally, H*γ*D-crys, akin to other proteins prone to aggregation, harbors a hydrophobic core within its structure, primarily composed of six amino acid residues.^5,6,15,16^ Among these residues, Met43, Phe56, and Ile81 belong to the N-terminal domain, while Val132, Leu145, and Val170 are part of the C-terminal domain. Mutations of these residues, particularly those within the N-td, have been experimentally demonstrated to significantly destabilize the protein structure and enhance its unfolding kinetics by facilitating access to new metastable states.^9,16^ To preserve the hydrophobicity of the interdomain interface, the wild-type H*γ*D-crys employs two flanking pairs, namely Gln54-Gln143 and Arg79-Met147, which act as molecular tethers, encircling the hydrophobic core formed by the N- and C-terminal domains.^15^ Moreover, its N- and C-terminals shield its hydrophobic core, with the C-terminal end forming a loop over the protein’s interdomain interface, resembling a structural anchor between the two protein lobes.

It is necessary to note that conformational perturbations that lead to cataract are generally subtle, as several cataract-associated single and double amino-acid substitutions within the wild type H*γ*D-crys have been previously reported to retain a native-like conformation showing no evidence of large scale misfolding.^7,11^ It therefore appears that crystallin aggregation is triggered by subtle conformational changes that may be difficult to capture experimentally. In contrast, environmental perturbations such as changes in temperature, or chemical denaturants, including urea and guanidine hydrochloride (GdnHCl), induce complete unfolding of the protein, henceforth significantly distorting its equilibrium conformational landscape.^15,16^

To bridge the gap between extensive conformational changes triggered by “harsh” stress factors and subtle conformational instabilities in aggregation-prone mutants, we turn to high hydrostatic pressure as a milder physical perturbation to protein structure. Crucially, nature favors life at extremely high pressure, as evidenced by genetic adaptations enabling deep-sea proteins to function at extreme pressures.^27^ Notably, even minor amino acid substitutions at critical functional sites can adapt the pressure sensitivity of a protein by favoring chemical conformations that are less prone to changes in volume.^28^ However, such adaptations are pressure-specific, limiting the depth at which deep-sea organisms can thrive while preserving *in vitro* sensitivity to pressure.^29^ Pressure affects protein folding through stabilizing folding intermediates with smaller specific volumes.^30–32^ Additionally, pressure serves as a valuable tool for inferring mechanistic information about protein folding and refolding by perturbing a protein’s thermodynamic equilibrium.^33–35^ At a molecular level, the response of proteins to high pressure predominantly involves changes to intramolecular hydrophobic interactions. However, it has been experimentally observed that increasing pressure can elicit both increases and decreases in protein stability.^36^ These intricacies have resulted in a resurgent interest in understanding the impact of high pressure on protein folding in general, and have further motivated our investigation of how pressure impacts the stability indicators associated with the N-terminal domain and the interdomain interface of H*γ*D-crystallin. To the best of our knowledge, this is the first computational investigation of how pressure impacts the conformational landscape of a large globular protein.

However, studying these subtle conformational changes within H*γ*D-crystallin poses a challenge for conventional experimental techniques due to their limited spatiotemporal resolution in capturing small variations in protein structure. Meanwhile, traditional computational methods such as Md might become inefficient in sampling conformational landscapes characterized by basins separated by substantial free energy barriers.^37,38^ To address these limitations, a wide variety of advanced sampling techniques have been developed to facilitate traversing energy barriers in Md simulations.^39–41^ Notably, replica exchange molecular dynamics (Remd) has emerged as a powerful tool for exploring rough potential energy landscapes with moderate bottlenecks, making it ideal for studying conformational landscapes of proteins.^42^ Here, we utilize Remd simulations to investigate the impact of pressure on the conformational landscape of the wild-type H*γ*D-crystallin, which we probe using principal component analysis (Pca). Our investigation quantifies the impact of pressure on both conventional global metrics of protein stability and localized structural features confirmed or hypothesized to play crucial roles in stabilizing the aromatic contacts within the N-terminal domain and modulating the hydration status of the hydrophobic interdomain interface. Intriguingly, our findings reveal correlations between the two primary Pca components and the equilibrium distributions of the local order parameters associated with the N-td and the interdomain interface. These observed correlations shed light on the relationship between conformational dynamics and key stability markers affected by pressure in H*γ*D-crystallin.

## II. METHODS

### A. Molecular dynamics simulations

All Md simulations are performed using the open-source Gromacs^43^ software, with Newton’s equations of motion integrated using the leapfrog algorithm with a time step of 2 fs. A modified Amber force-field is used to model the protein (Amber ff99SB-ILDN),^44^ while water molecules are represented using the Tip4p model.^45^ The reliability of this force field with Tip4p water model has been validated by other simulation studies on proteins.^46–48^ (We also confirm the stability of H*γ*D-crys upon utilizing this force-field, as depicted in Fig. S1.) All bond and angle constraints are maintained using the linear constraint solver (Lincs) algorithm.^49^ All production runs are conducted within the isothermal isobaric (Npt) ensemble with temperature and pressure controlled using the Nosé-Hoover thermostat^50^ and the Parinello-Rahman barostat,^51^ with time constants of 0.4 ps and 2 ps, respectively. All long-range electrostatic interactions are treated using the smooth particle mesh Ewald^52^ (Pme) method with a grid spacing of 0.1 nm, while all shortrange interactions are truncated at 1 nm.

### B. System setup and equilibration

The initial configuration of the wild-type human *γ*D-crystallin is obtained from the Research Collaboratory for Structural Bioinformatics (Rcsb) Protein Data Bank (Pdb code 1HK0).^7^ Subsequently, the electro-neutral protein is solvated with 15,133 water molecules, and the resulting system undergoes energy minimization employing 5,000 steps of the steepest descent method in order to remove any unphysical atomic overlaps. Following energy minimization, the system undergoes a two-step equilibration process. First, a 100-ps position-restrained equilibration within the canonical (Nvt) ensemble is conducted employing the Brendsen thermostat^53^ in order to properly equilibrate the solvent molecules surrounding the protein. Subsequently, a 100-ps Npt simulation is performed using the Parinello-Rahman barostat to stabilize density and pressure. After equilibration, a pressure ramp is applied starting from the Npt-equilibrated structures. Subsequently, 100-ns production runs are initiated at each target pressure, namely 1 bar, and 5, 7, 10, and 15 kbar, respectively.

### C. Dimensionality reduction

We employ principal component analysis^54^ (Pca) to analyze all our Md trajectories. Functioning as a linear dimensionality reduction technique, Pca identifies key collective variables pivotal in explaining the variability in data. Starting from a trajectory, *x*(*t*), which could be a collection of time frames, or samples taken from a probability distribution, the covariance matrix *C* is computed as,

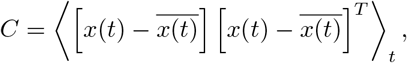

and is subsequently diagonalized. The eigenvectors of *C* corresponding to its leading eigenvalues serve as representative collective variables capturing substantial deviations from the mean, *x*(*t*). Pca has become a standard technique for constructing free energy landscapes of proteins.^55^ In our analysis, the trajectories fed into Pca consist of atomic coordinates of *α*-carbons (C*α*) of all residues. After conducting Pca, the fraction of variance explained (Fve), evaluating the proportion of variance retained by each eigenvalue is computed as,

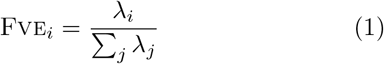

The cumulative Fve for a set of eigenvalues can be determined by adding up their respective individual Fves. To investigate the conformational landscape at different pressures, we project 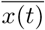onto the two primary eigenvectors of *C*. These individual projections are then used for constructing free energy profiles as discussed in the text.

### D. Replica exchange Md simulations

Replica exchange Md constitutes an advanced sampling technique rooted in the idea of parallel tempering, and was initially proposed by Sugita and Okamoto in 1999.^40^ This method enables effective exploration of rugged potential energy landscapes by concurrently conducting multiple simulations at varying temperatures and intermittently exchanging configurations between adjacent simulation “windows” based on their respective Boltzmann weights. In this work, we conduct two rounds of Remd simulations, each employing 35 replicas within the temperature range, 290 K ≤ *T* ≤ 342 K. The temperature differences between adjacent replicas are carefully selected to maintain an exchange acceptance rate of approximately 20− 25%. The initial round of simulations, spanning 90 ns, commences with configurations periodically collected along the 100-ns conventional Md trajectories outlined in Section II B. However, due to the rugged nature of the potential energy landscape at elevated pressures, effectively exploring the free energy landscape, particularly the emerging metastable basins, poses a challenge even with extended trajectories. To address this challenge, we randomly select 35 configurations from the two Pca basins identified at 5 kbar with equal probabilities. These configurations serve as initial points for a subsequent 60-ns round of simulations. For ambient pressure, configurations are randomly chosen from those periodically collected at ambient temperature during the initial round of Remd. All analyses and results presented in this manuscript rely on trajectories obtained during this second round of Remd. We ensure the adequacy of sampling and therefore the robustness and reliability of our findings, by verifying that extending the Remd trajectories does not alter the qualitative shapes or positions of Pca basins.

### E. Structural analysis

In order to assess how H*γ*D-crys stability is impacted by environmental perturbations, a collection of global and local order parameters (Ops) are computed. The global OPs computed here are all standard measures of protein stability. The first is the *root mean square deviation (RMSD)*, expressed as:

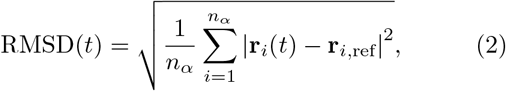

The summation is performed across all *n*_*α*_ C*α* atoms. **r**_*i*_(*t*) represents the instantaneous position of the *i*th C*α* at time *t*, while **r**_*i*,ref_ denotes its corresponding position within the protein’s least-squares fitted reference structure, i.e., the solvated Rcsb structure.Another important metric for assessing protein stability is the *radius of gyration, R*_*g*_, defined as,

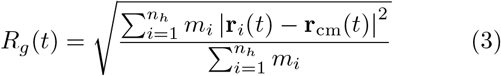

Here, *m*_*i*_ and **r**_*i*_(*t*) correspond to the mass and instantaneous position of atom *i*, respectively. The summation is over all *n*_*h*_ heavy (non-hydrogen) atoms within the protein. This approach is chosen to minimize the influence of highly fluctuating hydrogen nuclei on the computed *R*_*g*_.

In addition to RMSD and radius of gyration, we also compute the *solvent-accessible surface area (SASA)* using the double cubic lattice method.^56^ This also enables us to compute the average volume of the protein, more precisely van der Waals volume approximation of Connolly molecular volume,^57^ using the Gauss-Ostrogradskii theorem. The change in protein’s volume Δ*V* (with respect to ambient conditions) is then computed using,

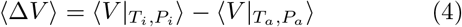

Here, the subscripts *i* and *a* refer to target and ambient conditions, respectively.

The local OPs considered in this work are all specific to H*γ*D-crys and correspond to distances between specific residues, as will be discussed in Section III. Unless otherwise specified, the distance between two residues is defined as the minimum distance between their heavy atoms. More precisely, if residues *i* and *j* are comprised of *n*_*i*_ and *n*_*j*_ heavy atoms, the instantaneous distance between them is defined as,

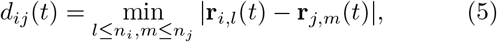

where **r**_*i,l*_ and **r**_*j,m*_ correspond to the position of the *l*-th and *m*-th heavy atom within the *i*-th and *j*-th residue, respectively.

## III. RESULTS AND DISCUSSION

### A. Global order parameters

In order to assess the impact of pressure and temperature on the native structure of H*γ*D-crys, we first calculate the global Ops described in Section II E, with their averages and error bars depicted in Fig. 2. Since elevating pressure results in an increase in the density of the water molecules surrounding the protein, both ⟨RMSD⟩ (Fig. 2A) and ⟨*R*_*g*_*⟩* (Fig. 2B) tend to decrease with pressure, indicating enhanced compactness and stiffness of the protein upon compression. In contrast, increasing temperature results in an increase in both these quantities (Figs. 2A,B). A similar trend is observed for ⟨SASA⟩, which increases slightly with temperature (Fig. 2C) but exhibits a reduction of approximately 3 nm^2^ with pressure (Fig. 2C). This observation suggests that exposing the protein to high pressure has a more pronounced impact on its overall hydration structure compared to elevating temperature. This further reinforces the contrasting effects of temperature and pressure on these global order parameters.

**FIG. 2.**
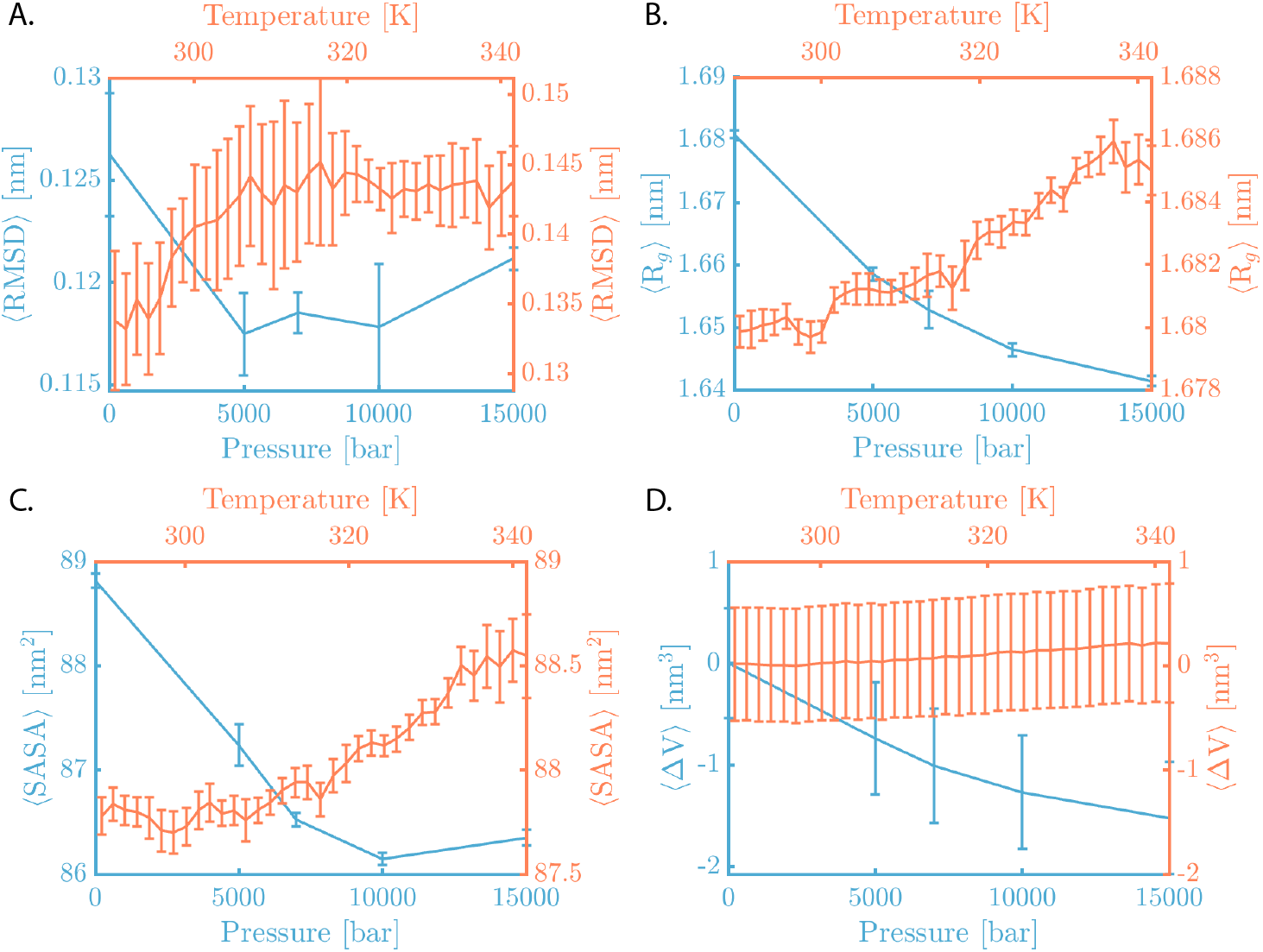
Dependence of global measures of protein stability to temperature and pressure: (A) root mean-square deviation, (B) radius of gyration, (C) solvent accessible surface area, and (D) change in protein volume.

Fig. 2D depicts the change in protein volume upon increasing pressure and temperature. Once more, the influence of high pressure on ⟨ΔV⟩ is directionally different from that induced by high temperature. Additionally, akin to SASA, ⟨ΔV⟩ undergoes more substantial changes with pressure than temperature. The impact of pressure on ⟨ΔV⟩ aligns with Le Chatelier’s principle, which predicts that an increase in pressure will prompt a shift in thermodynamic equilibrium towards configurations with smaller volumes.

The slight changes observed in these global measures of protein stability upon increasing temperature and pressure align well with the known high stability of H*γ*D-crys, which is highly resistant to unfolding. These global order parameters typically increase with temperature, yet the extent of this increase remains notably minimal within the considered temperature range. Conversely, in response to pressure, they uniformly exhibit a slight decrease, indicating that pressure-induced conformational changes are likely local in nature, while the overall fold remains mostly unchanged. This is consistent with earlier investigations of pressure-induced unfolding, which generally culminate in the formation of more compact conformations.^35^

### B. Free energy landscapes

Our examination of global order parameters confirms that the impact of pressure on the global structure of H*γ*D-crystallin remains relatively mild, even under pressures as large as 15 kbar. This implies that increasing pressure only induces subtle changes in the protein’s structure, evading detection through global order parameters. To identify and characterize such changes, we first scrutinize our Remd trajectories using Pca. However, the efficacy of Pca hinges upon the protein’s local dynamics possessing sufficient simplicity to be amenable to the linearization inherent in Pca. Under such conditions, crucial conformational changes within the protein can be succinctly captured through a limited number of principal components. We believe that the absence of dramatic changes in H*γ*D-crys conformation, and the overall stability of global Ops justify the use of Pca here.

Based on the Fves computed across all target pressures (Fig. 3A), the first two leading eigenvalues capture over 50% of variability in data. Consequently, we project each Remd trajectory onto the corresponding eigenvectors associated with these leading eigenvalues, resulting in a time series projection, (*γ*_1_(*t*), *γ*_2_(*t*)), with *γ*_*i*_ corresponding to the projection along the *i*th principal component. Subsequently, Δ*G*(*γ*_1_, *γ*_2_) is computed as follows. First, the (*γ*_1_, *γ*_2_) space is discretized into distinct bins, and the number of observations within each bin (𝒩_*ij*_) is determined. The free energy associated with each bin is then estimated as:

**FIG. 3.**
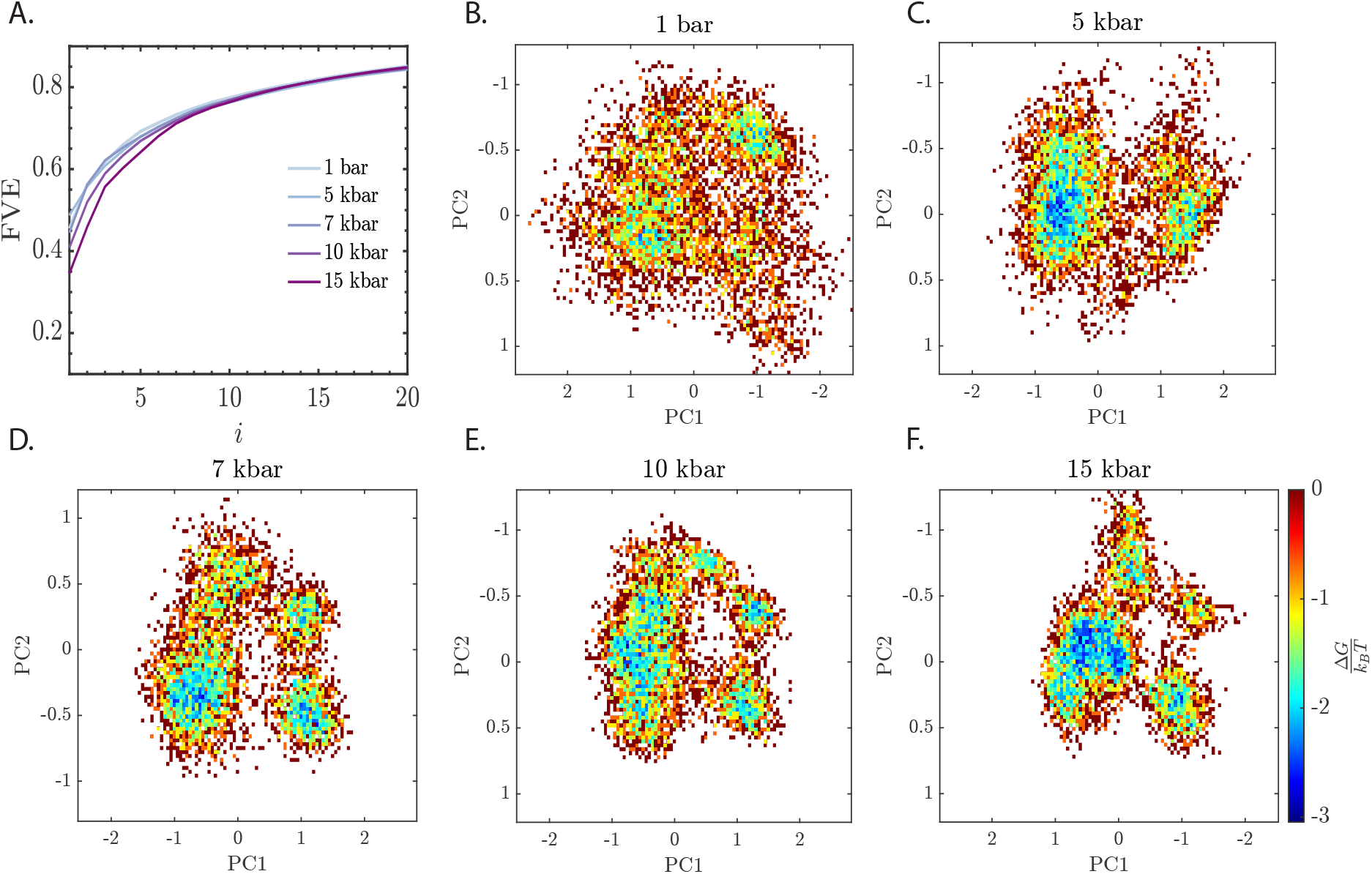
(A) Cumulative fraction of variance explained (Fve) for the first twenty eigenvalues of all target pressures. (B-F) Free energy profiles within the space of projections along the two principal components at (B) 1 bar, (C) 5 kbar, (D) 7 kbar, (E) 10 kbar and (F) 15 kbar.

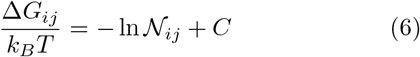

where *k*_*B*_ denotes the Boltzmann constant and *C* is a normalization constant.

It is clearly evident that increasing pressure results in the emergence of new basins within the free energy landscape. Moreover, such basins are separated by modest free energy barriers of approximately 2− 3*k*_*B*_*T*. This is in contrast to ambient pressure wherein Pca projections scatter within a single free energy basin, comprised of shallow wells separated by barriers of approximately 0.5*k*_*B*_*T*. The structuring of the free energy landscape at elevated pressures suggests the preferential stabilization of specific conformational states. To discern and characterize such conformational states and to associate them with different basins within the free energy landscape, it is imperative to identify local physically interpretable order parameters that correlate with Pca basins. In search of such local Ops, we focus on structural features that have been experimentally demonstrated to correlate with denaturation under environmental perturbations. Specifically, we investigate aromatic contacts within the Nterminal domain, as well as markers associated with the stability and hydration of the hydrophobic core.

### C. Local order parameters

#### 1. N-terminal domain stability

As detailed in Section I, aromatic contacts within the Ntd have been demonstrated to be critical for H*γ*D-cryst stability (Fig. 1A).^5,15–17,58^ We therefore probe the impact of pressure on the minimum distances among specific aromatic pairs known to establish contacts within both the N-td and C-td. Specifically, we examine six tyrosine-tyrosine pairs (Y16-Y28, Y45-Y50, Y45-Y62, Y55-Y62, Y92-Y-97, and Y133-138), two tyrosine-tryptophan pairs (Y55-W68, Y62-W68), one tyrosine-phenylalanine pair (Y6-F11), and one phenylalanine-phenylalanine pair (F115-F117). Among these aromatic pairs, seven are associated with the reportedly less stable N-td, while the remaining three pairs facilitate interactions within the C-td.

Figure 4 illustrates the evolution of the histograms of these ten minimum distances upon increasing pressure. In general, most of these distances tend to decrease as pressure increases, with only a few remaining almost unchanged, such as Y6-F11 (Fig. 4A), Y45-Y50 (Fig. 4C) and Y92-Y97 (Fig. 4H). For a few pairs, however, the qualitative shapes of the histograms also change in addition to leftward shifts in their means. Specifically, both Tyr45-Tyr62 (Fig. 4D) and Tyr55-Tyr62 (Fig. 4E) pairs reveal the emergence of new peaks at 1.12 nm (Fig. 4D) and 0.65 nm (Fig. 4E), respectively. Furthermore, the singular peak of the Tyr16-Tyr28 distance bifurcates into two distinct peaks under high-pressure conditions (Fig. 4B).

**FIG. 4.**
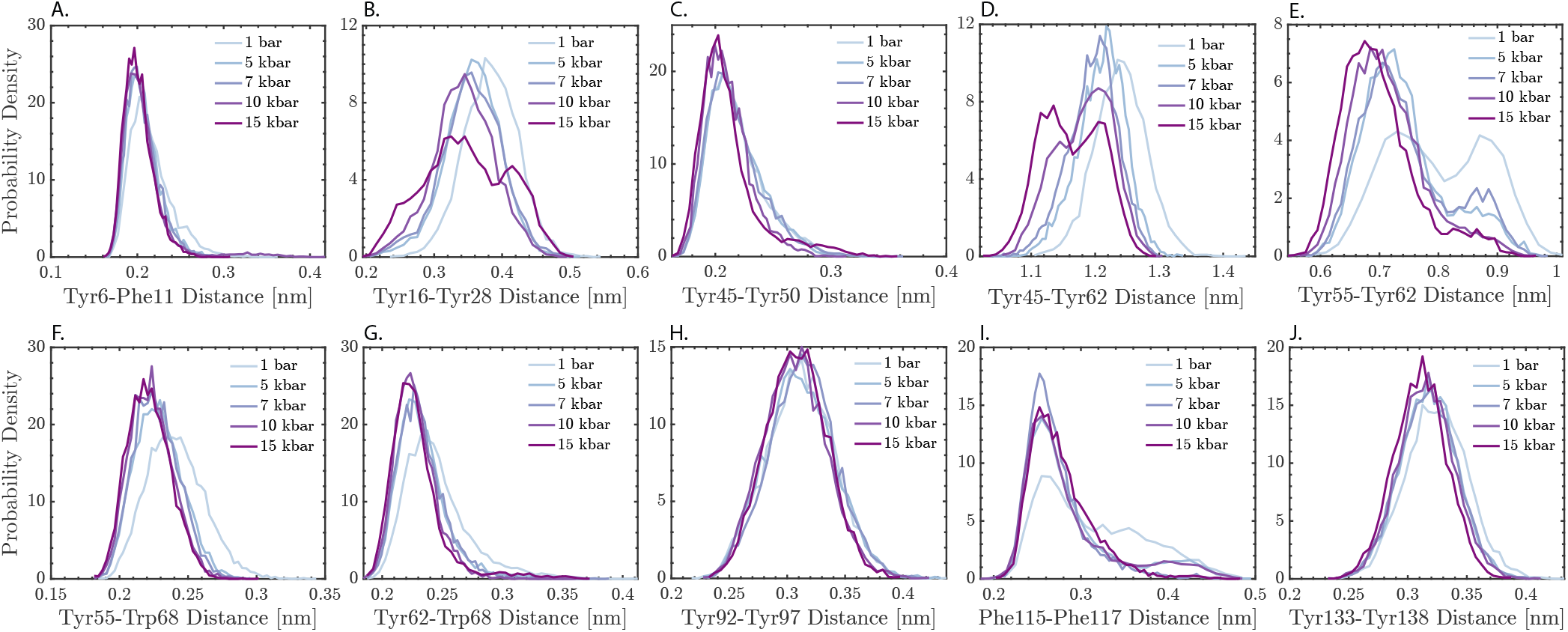
Histograms of minimum distances of the ten aromatic pairs previously identified as potential markers of H*γ*D-crys stability as a function of pressure. Among these (A-G) and (H-J) correspond to contacts within the N-td and C-td, respectively.

Based on these observations, it is reasonable to surmise the existence of a potential correlation between (some of) these distances and the basins identified through Pca. To facilitate the visual exploration of such possibility, we implement an agglomerate hierarchical cluster linkage algorithm to systematically group Pca projections into clusters, color-coding them based on their association with different principal component projections. Specifically, clusters linked to distinct values of PC1 are denoted in purple and green, while those associated with distinct PC2 values are represented by darker and lighter shades of each color (Figs. 5A-D). Subsequently, conditional histograms of the minimum distances considered in Fig. 4 are constructed to examine the existence of any correlations between either of those distances and the first two Pca projections.

**FIG. 5.**
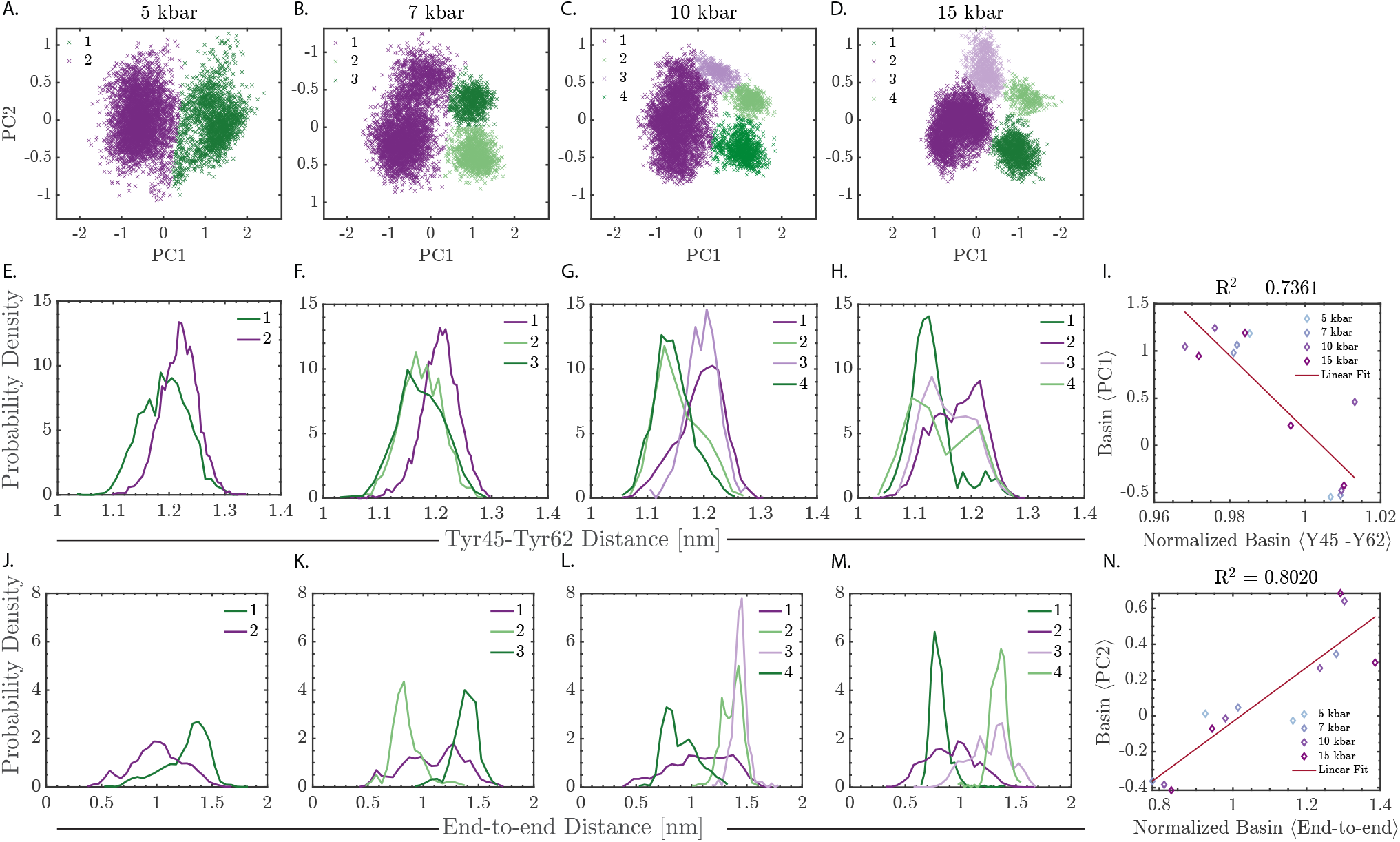
(A-D) Distinct free energy basins within the PC1-PC2 space colored based on their association with different principal component projections: green and purple for PC1 and dark and light for PC2. (E-H, J-M) Conditional histograms of (E-H) Y45-Y62 and (J-M) end-to-end minimum distances for the basins distinctly colored in (A-D). (I, N) Linear correlations between the normalized mean (I) Y45-Y62 and (N) end-to-end distances with (I) mean PC1 and (N) mean PC2 per basin, respectively.

Among all the pairs considered in Fig. 4, the Tyr45Tyr62 pair (Fig. 1B) exhibits the strongest correlation with the first principal component projection (PC1), with the other pairs exhibiting no meaningful correlation (Figure S2). As depicted Fig. 4D, the Y45-Y62 minimum distance histogram exhibits a peak at 1.25 nm at ambient pressure. Upon increasing pressure, this peak undergoes a leftward shift. Moreover, a second peak emerges at approximately 1.15 nm, initially appearing faintly at 5 kbar (Fig. 5E), but gaining prominence at 7 kbar (Fig. 5F) and beyond. Examining the conditional histograms of the Tyr45-Tyr62 distance (depicted in Figs. 5E-H) elucidates a discernible correlation with PC1. Specifically, basins associated with smaller PC1 values consistently exhibit reduced Y45-Y62 distances, despite some overlaps evident in the respective histograms.

In order to make our analysis more quantitative, we compute the mean values of PC1 and the Y45-Y62 distance for each Pca basin. Moreover considering the concurrent decrease in the mean Y45-Y62 distance with pressure, we normalize each mean basin distance by the total mean distance at the corresponding pressure. As illustrated in Fig. 5I, a meaningful correlation emerges between the normalized basin mean Y45-Y62 distances and PC1 values, displaying a correlation coefficient of *R*^2^ = 0.7361. This correlation underscores the association between the leading principal component projections and the structural alterations signified by a change in the Y45-Y62 distance.

The situation is more nuanced for other top contenders, most notably the Tyr55-Tyr62 pair (Figs. S2JN). At ambient pressure, the minimum distance histogram exhibits two peaks at 0.73 nm and 0.89 nm (Fig. 4B). However, increasing pressure leads to a distinct leftward shift of the first peak, accompanied by a gradual reduction in the prominence of the second peak, which starts diminishing at 5 kbar and nearly disappears by 10 kbar. Despite the evident pressure-induced changes in the shape of the histogram, we fail to discern any significant correlation between the absolute or normalized mean distances within the Pca basins and either of the mean Pca projections (Fig. S2N). A similar scenario unfolds for other important contenders, such as Y16-Y28 (Figs. S2E-I) and F115-F117 (Figs. S2O-S), where changes in histogram shapes occur with increasing pressure, yet meaningful correlations with Pca projections are not apparent (Figs. S2I,S).

Collectively, these observations suggest that pressure stabilizes the native contacts among the aromatic residues, most importantly the tyrosine pairs, thereby intensifying the crowding effect within the *β*-sheet motifs of the N-td domain. This phenomenon aligns with observations from studies on proteins of comparable sizes, where the crowding of *β* sheets has been identified as a hallmark of heightened native structural stability.^21–23,59^ Moreover, our observations suggest a correlation between the strength of aromatic contacts within the N-terminal domain and the first principal component from our Pca analysis.

#### 2 Interdomain interface stability

Another pivotal determinant of H*γ*D-crys stability is the integrity of the interdomain interface, which, if compromised due to mutations or environmental variations, can adversely affect the stability of the hydrophobic core within the protein.^5,6,15,16^ To investigate the pressure-induced effects on the stability of interdomain interface, we examine several associated local order parameters.

The initial set of such order parameters focus on quantifying the proximity between the Nand C-terminals. In the H*γ*D-crys structure, the C-terminal extends over the interdomain interface, serving as both an anchoring point between protein lobes and a barrier to the protein’s hydrophobic core. In particular, we consider the following order parameters: (i) the minimum distance between the first residue of the N-terminal and the last residue of the C-terminal, representing the initial N-td and the terminal C-td residues, respectively, (ii) the distance between the centers of mass of the structureless loops of the N- and C-terminals, and (iii) the distance between the centers of mass of residues 80-81 and 168-170, corresponding to the closest *β*-sheets of the interdomain interface. The histograms of these order parameters as a function of pressure are depicted in Figs. 6A-C. Unlike the consistent decrease observed in aromatic pair distances with increasing pressure, the influence of pressure on these order parameters is more intricate and therefore less straightforward to interpret. The center of mass distances between the N- and C-terminals, and the two closest *β*-sheets, generally exhibit a decline with escalating pressure (Figs. 6B,C). The end-to-end distance, however, demonstrates a rich and intricate behavior (Fig. 6A). Despite its mean being virtually insensitive to pressure, the positions and magnitudes of peaks within its histogram exhibit a complex dependence on pressure.

**FIG. 6.**
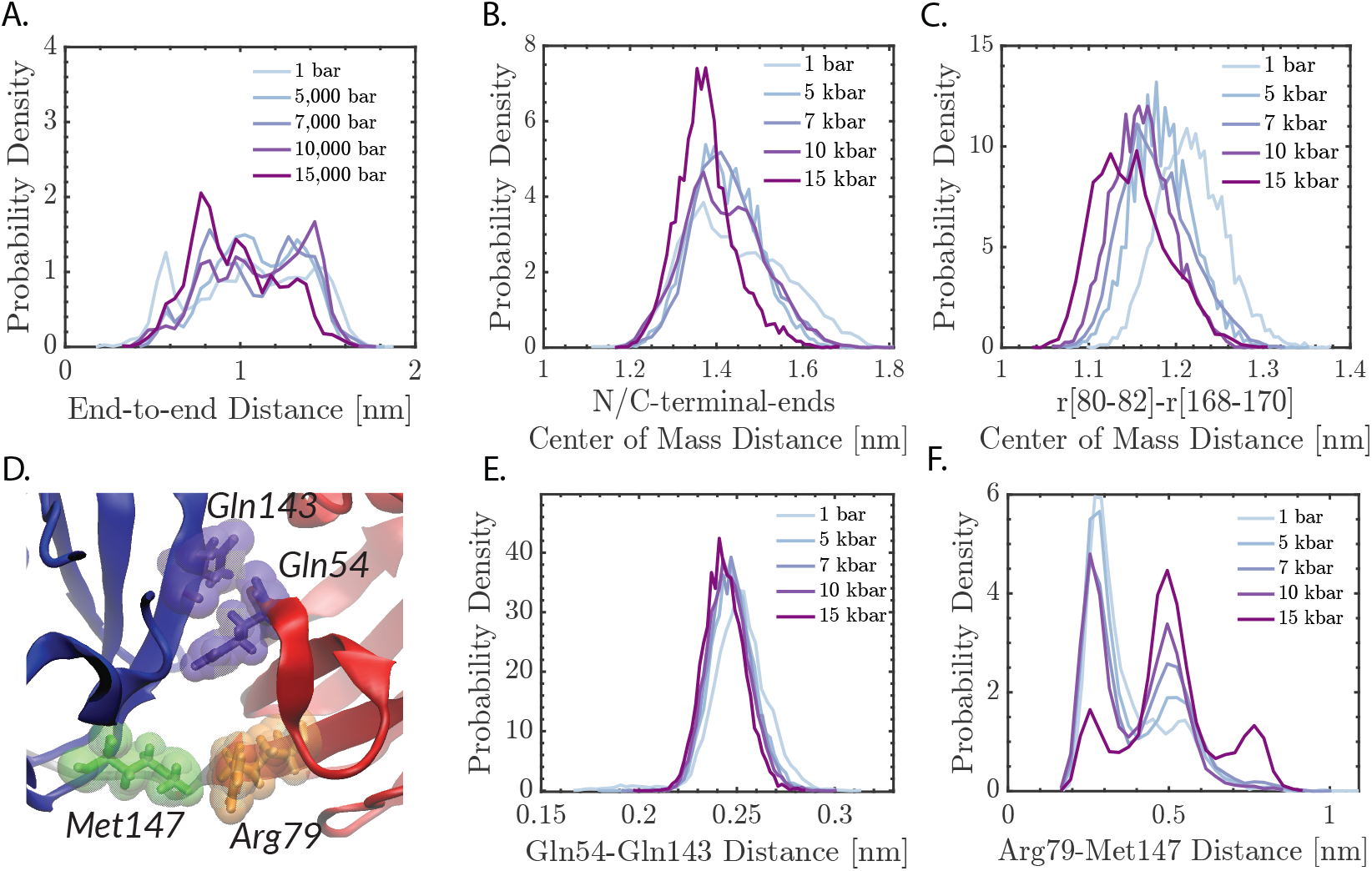
Histograms of local order parameters associated with the interdomain interface of H*γ*D-crys, namely distance between (A) N- and C-terminal residues, (B-C) centers of masses of structureless regions within each terminal and (C) residues 80-82 from the N-terminal and residues 168-170 from the C-terminal. (D) Rendering of H*γ*D-crys highlighting the flanking pairs Gln54-Gln143 and Arg79-Met147, (E-F) Histograms of minimum distances between (E) Gln54-Gln143 and (F) Arg79-Met147 as a function of pressure.

The second category of order parameters are associated with the Gln54-Gln143 and Arg79-Met147 pairs, the two flanking pairs highlighted in Fig. 6D and discussed in Section I that function as gateways to the hydrophobic core.^15^ The histograms of minimum distances of both pairs are depicted in Figs. 6E and F. As depicted in Fig. 6E, pressure has minimal impact on the Gln54-Gln143 pair, leaving their distance virtually unchanged. In contrast, the histogram corresponding to the Arg79-Met147 pair distance (Fig. 7F) exhibits a shift towards larger values at higher pressures. Once more, this underscores that the effect of pressure on the interdomain interface is complex and nontrivial, which is in stark contrast to the behavior observed for aromatic contacts.

**FIG. 7.**
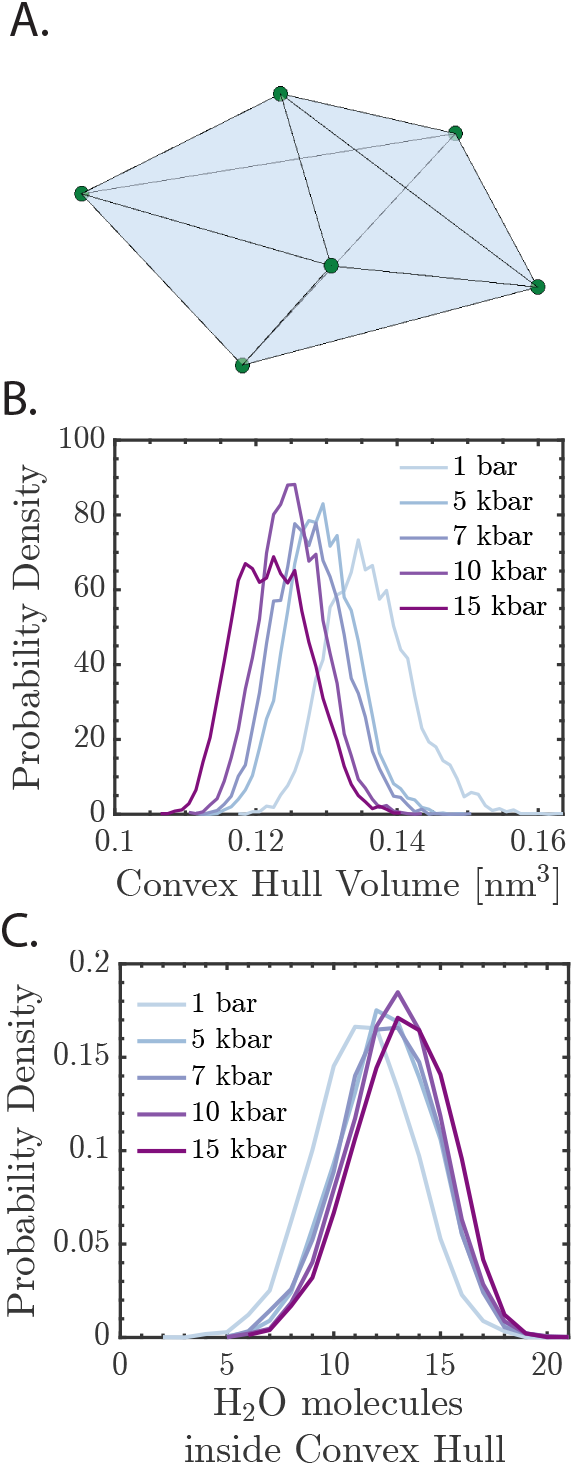
(A) Schematic representation of the convex hull approximation of the hydrophobic core using the centers of masses of the six hydrophobic residues, Met43, Phe56, Ile81, Val132, Leu145, and Val170 (shown in Fig. 1C). (B-C) Histograms of (B) the volume of and (C) the number of water molecules within the convex hull as a function of pressure.

Upon closer examination of conditional histograms of these order parameters for basins identified from Pca, we only find a distinct separation between the conditional histograms of the end-to-end distance histograms (Fig. 6A) associated with basins of distinct PC2 values. Naturally, such a distinction is absent at 5 kbar where both basins share the same PC2 values (Fig. 5J) and manifests itself at 7 kbar (Fig. 5K) and above (Figs. 5L,M) where basins with distinct PC2 values emerge. Similar to the approach discussed in Section III C 1, we compute the mean end-to-end distance per basin and normalize it with the corresponding mean at each pressure. As illustrated in Fig. 5N, we observe a strong correlation (with a correlation coefficient *R*^2^ = 0.8020) among these normalized distances and the corresponding per-basin mean PC2 values. A weaker, but nonetheless meaningful, correlation (*R*^2^ = 0.7153) is observed between mean center-of-mass distance of the loop regions of Nand C-terminals (Fig. S3I). This is not surprising considering the expected correlation between that center of mass distance and the end-to-end distance. No strong correlation was observed for the other three order parameters considered in Fig. 6 (see Fig. S3).

This observation confirms that the two collective variables derived from Pca are linked to physically interpretable local order parameters. More specifically, the Y45-Y62 aromatic pair contact within the N-terminal domain correlates with PC1, while the end-to-end distance correlates to PC2. Consequently, the basins identified via Pca do not represent modeling artifacts; rather, they possess distinct and clear physical interpretations.

Motivated by the correlation between the second principal component and the end-to-end distance, which is a proxy for the stability of the interdomain interface, we probe the effect of pressure on the size and the solvent exposure of the hydrophobic core. To quantify the size of the hydrophobic core, we employ the volume of the convex hull (Fig. 7A) constructed from the centers of masses of six crucial hydrophobic residues pivotal for maintaining the integrity of the interdomain interface. As expected, the volume of the hydrophobic core decreases upon increasing pressure, as indicated by the histograms of convex hull volumes presented in Fig. 7B. Interestingly though, the number of water molecules enclosed within the convex hull rises with pressure (Fig.7C), implying that pressure alters water’s density and structure around the protein in a manner that counteracts the reduction in the accessible volume of the hydrophobic core. Evaluating the number of hydrogen bond donor-acceptor pairs surrounding the six hydrophobic residues reveals a minor increment in the number of protein-solvent hydrogen bonds, while the number of intramolecular hydrogen bonds within the hydrophobic core remains mostly unchanged (Fig. S4). This observation once again validates the intrusion of water into the hydrophobic core under high-pressure conditions. The expansion of the hydrophobic core is likely triggered by the increase in the Arg79-Met147 pair distance as depicted in Fig. 7F.

## IV. CONCLUSIONS

In this study, we utilize Remd simulations augmented with Pca to explore the conformational landscape of human *γ*D-crystallin under elevated pressures. As anticipated, our findings reveal that high pressure only has a modest impact on global measures of protein stability, such as RMSD and radius of gyration. However, through the projection of Remd trajectories using the first two principal components, an intriguing picture emerges: the singular basin observed at ambient pressure divides into multiple distinct basins with increasing pressure. We probe multiple local order parameters previously demonstrated or hypothesized to be markers of H*γ*D-crys stability, namely various aromatic contacts within the N-td, and a handful of characteristic distances associated with the interdomain interface. Our analyses reveals a correlation between the minimum distance of the aromatic pair, Tyr45-Tyr62, with PC1, as well as a relationship between the N-/C-td end-to-end distance and PC2. Furthermore, we observe the contraction of the hydrophobic core, accompanied by its intrusion by water molecules, possibly facilitated by the increased separation between Arg79 and Met147. These findings collectively provide deeper insights into the structural alterations occurring within H*γ*D-crystallin under elevated pressure conditions, shedding light on specific molecular changes influencing its stability.

It is generally understood that pressure affects protein folding through favoring intermediates that poses smaller specific volumes, which can result in both stabilizing and destabilizing the folded state. When it comes to H*γ*Dcrystallin, it is unclear whether elevated pressures lead to its stabilization or disruption. On one hand, the consistent reduction in distances between aromatic pairs within the N-td suggests a potential overall stabilizing effect. Conversely, the infiltration of water molecules into the hydrophobic core implies a potential destabilization of the interdomain interface. Consequently, the overall effect of pressure on the stability of H*γ*D-crystallin remains undetermined, as the opposing influences on different structural elements necessitate a comprehensive evaluation to ascertain the net impact.

To comprehensively address this issue, it is imperative to determine the change in folding free energy, typically achieved through methods such as thermodynamic integration that rely on the two-state model for protein folding.^35,60^ However, this task poses significant challenges due to substantial free energy barriers separating the folded and unfolded states of large proteins, such as H*γ*D-crys. Consequently, standard enhanced sampling techniques, such as Remd, struggle to efficiently capture unfolding-folding transitions despite their ability to sample rugged free energy landscapes. Remedying this issue might involve utilizing bias-based or path sampling techniques. However, such methods often require the specification of predefined reaction coordinates, which are generally unknown *a priori* for a complex process such as protein folding. Moreover, the accuracy of the two-state folding model remains unverified, contributing an additional layer of uncertainty to this conundrum. This uncertainty, coupled with the substantial energy barriers, renders the computation of folding free energy a daunting task that is beyond the scope of this work.

We also wish to emphasize that the pressure-induced conformational changes observed here can be characterized using a host of other dimensionality reduction techniques. Our choice of Pca is motivated by the subtle nature of the observed changes to H*γ*D-crystallin’s conformation. Through Pca, we establish meaningful correlations between crucial physically interpretable local order parameters and projections along the two primary principal components. However, it is important to note that alternative nonlinear methods, such as diffusion maps,^61,62^ isometric feature maps^63^ (Isomaps), and *t*-distributed stochastic neighbor embedding^64^ (*t*-SNE), might offer increased efficacy in capturing relevant low-dimensional embeddings of the free energy landscapes, potentially yielding a qualitatively different picture. Nonetheless, a significant challenge with applying such methods lies in the often less straightforward interpretability of the collective variables they produce. Assessing the sensitivity of our findings to the employed dimensionality reduction technique can be the focus of future explorations.

To identify physically interpretable collective variables that align with the leading principal component projections, we focus on local order parameters previously established or suggested as meaningful indicators of H*γ*Dcrystallin stability in prior experimental studies. While this targeted strategy enhances our ability to pinpoint relevant local order parameters, it also poses a potential limitation. By relying primarily on previously identified parameters, we may inadvertently restrict our exploration, potentially overlooking alternative local order parameters that might contribute to understanding the system but have not been detected in earlier investigations.

We wish to conclude with a few broader implications of investigating the conformational landscape of globular proteins such as H*γ*D-crystallin under elevated pressures. Examining pressure effects is relevant to the field of extreme biology, offering insights into the physiology of organisms adapted to high-pressure environments, such as the deep ocean or subsea floor.^27–29^ The adaptability of biological macromolecules to elevated pressures serves as a foundation for diverse biotechnological applications utilizing high pressure.^65^

Furthermore, the use of pressure, often combined with co-solvents, has emerged as a valuable tool in investigating the stability, energetics, packing, and intermolecular interactions of protein aggregates, such as amyloid fibrils.^66^ These fibrils result from the misfolding of proteins such as insulin, islet amyloid polypeptide, *β*2 microglobulin, amyloid *β*, and *α*-synuclein in various tissues.^67–69^ Specifically, high hydrostatic pressure facilitates distinguishing various stages of amyloid formation, enabling the acquisition of crucial structural and thermodynamic data at different transformation stages. This approach provides valuable insights into the mechanisms underlying amyloid formation and offers opportunities for developing therapeutic interventions targeting these protein misfolding disorders.

Another intriguing aspect of pressure effects lies in the striking resemblance between the structural changes in pure water under high pressure and those induced by the introduction of solutes. This observation suggests a correlation between such similarities and the subsequent dynamic processes. For instance, it is well established that the rate of homogeneous ice nucleation in both aqueous solutions and pressurized pure water depends solely on the water activity.^70^ Given the paramount importance of water structure and dynamics in protein folding, it is plausible to posit a similar connection between water activity and the thermodynamics as well as kinetics of protein folding. Consequently, investigating pressure effects becomes appealing as pressure could serve as a mild perturbation mimicking solute effects. This is important as under physiologically relevant conditions, nearly all globular proteins fold within aqueous solutions comprised of a large number of different solute types. Therefore, exploring pressure effects would offer an attractive alternative to model diverse solutes if the existence of such a hypothesized correlation can be established. This is an exciting possibility that can be the topic of future experimental and computational explorations.

On a related note, the impact of high pressure on H*γ*Dcrys is most vividly evident in the negative ⟨Δ*V⟩*, an effect that can also be induced by other perturbations such as crowding agents and co-solutes. For instance, negative ⟨Δ*V⟩* can arise from the disruption of salt bridges and ionic bonds, along with associated changes in hydration due to the presence of ionic salts. Considering these diverse influences, it becomes crucial to investigate the effect of co-solutes, including simple monovalent ions (such as K^+^) and small organic molecules, on both the conformational landscape of the monomer and its capacity to assemble into high molecular weight oligomers. Such research will contribute significantly to understanding the intricate interplay between various perturbations and their impact on the behavior and structural characteristics of H*γ*D-crys.

## ACKNOWLEDGMENTS

We thank J. Howard for useful discussions. A.H.-A. gratefully acknowledges the support of the National Science Foundation CAREER Award (CBET-1751971). These calculations were performed at the Yale Center for Research Computing (YCRC). This work used the Extreme Science and Engineering Discovery Environment (XSEDE), which is supported by National Science Foundation grant no. ACI-1548562.

