## Supplementary Information for "Effect of Pressure on the Conformational Landscape of Human *γ*D-crystallin from Replica Exchange Molecular Dynamics Simulations"

(Dated: January 6, 2024)

### S1. TEMPORAL EVOLUTION OF RMSD ALONG TWO ROUNDS OF REMD TRAJECTORIES

In order to ensure the suitability of the employed force-field for H$\gamma$D-crys, we monitor the time series of room-temperature RMSD along two REMD trajectories conducted at ambient pressure. Our analysis reveals consistent stability in the RMSD throughout both trajectories (Fig. S1), signifying that the folded conformation of H$\gamma$D-crys remains unchanged without undergoing disruptive alterations. We wish to reiterate that all analyses presented in the main text are based on the results obtained from the second round of REMD trajectories.

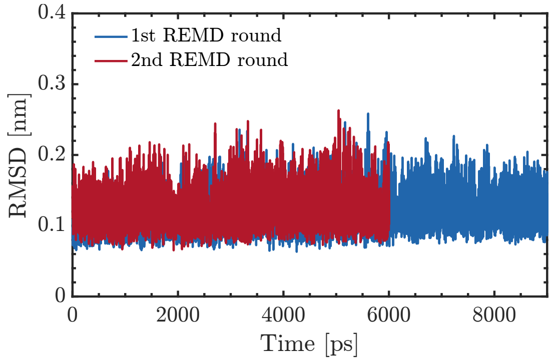


FIG. S1. Room-temperature ambient-pressure RMSD time series along the two rounds of REMD trajectories carried out for 90 ns (first round) and 60 ns (second round).

### S2. ANALYSIS OF CORRELATIONS BETWEEN LOCAL ORDER PARAMETERS AND PRINCIPAL COMPONENT PROJECTIONS

In Fig, 5, we present the two local order parameters that exhibit the strongest correlation with the leading principal component projections. To identify them, however, we conduct an extensive screening of several other potential order parameters. Here, we present our analysis of all promising candidates, i.e., order parameters that demonstrate intriguing pressure-dependent histograms, and yet exhibit a weak correlation with the principal component projections. Specifically, our examination encompasses Y16-Y28, Y55-Y62, and F115-F117 for PC1 (illustrated in Fig. S2). Additionally, for PC2 (depicted in Fig. S3), we scrutinize the N/C-terminal center of mass distance, the r[80-82]-r[168-170] center of mass distance, and the Arg79-Met147 minimum distance.

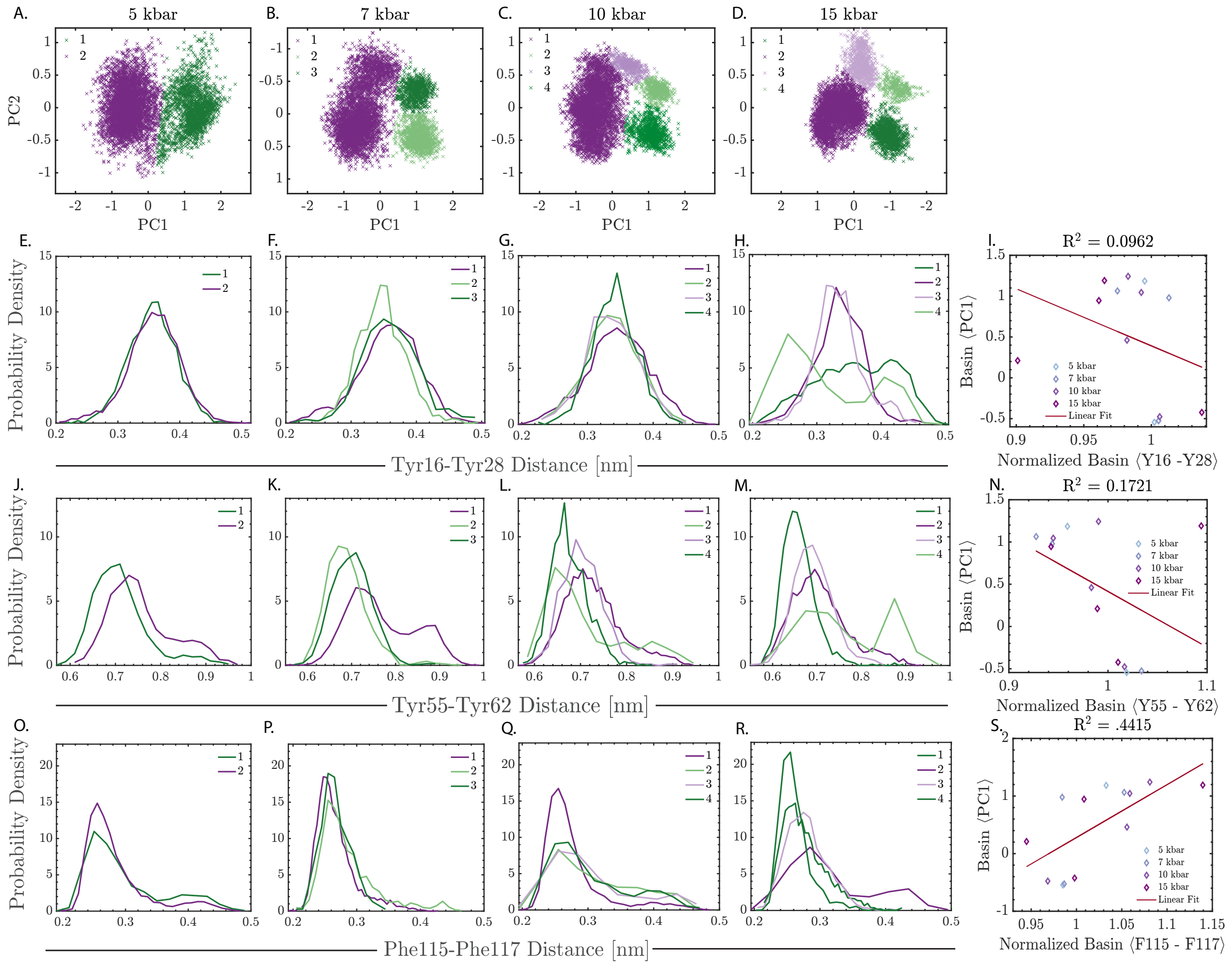


FIG. S2. (A-D) Distinct free energy basins within the PC1-PC2 space colored based on their association with different principal component projections: green and purple for PC1 and dark and light for PC2. (E-H, J-M, O-R) Conditional histograms of (E-H) Y16-Y28, (J-M) Y55-Y62, and (O-R) F115-F117 distances for the basins distinctly colored in (A-D). (I, N, S) Linear correlations between the normalized mean (I) Y16-Y28, (N) Y55-Y62, and (S) F115-F117 distances with mean PC1, respectively.

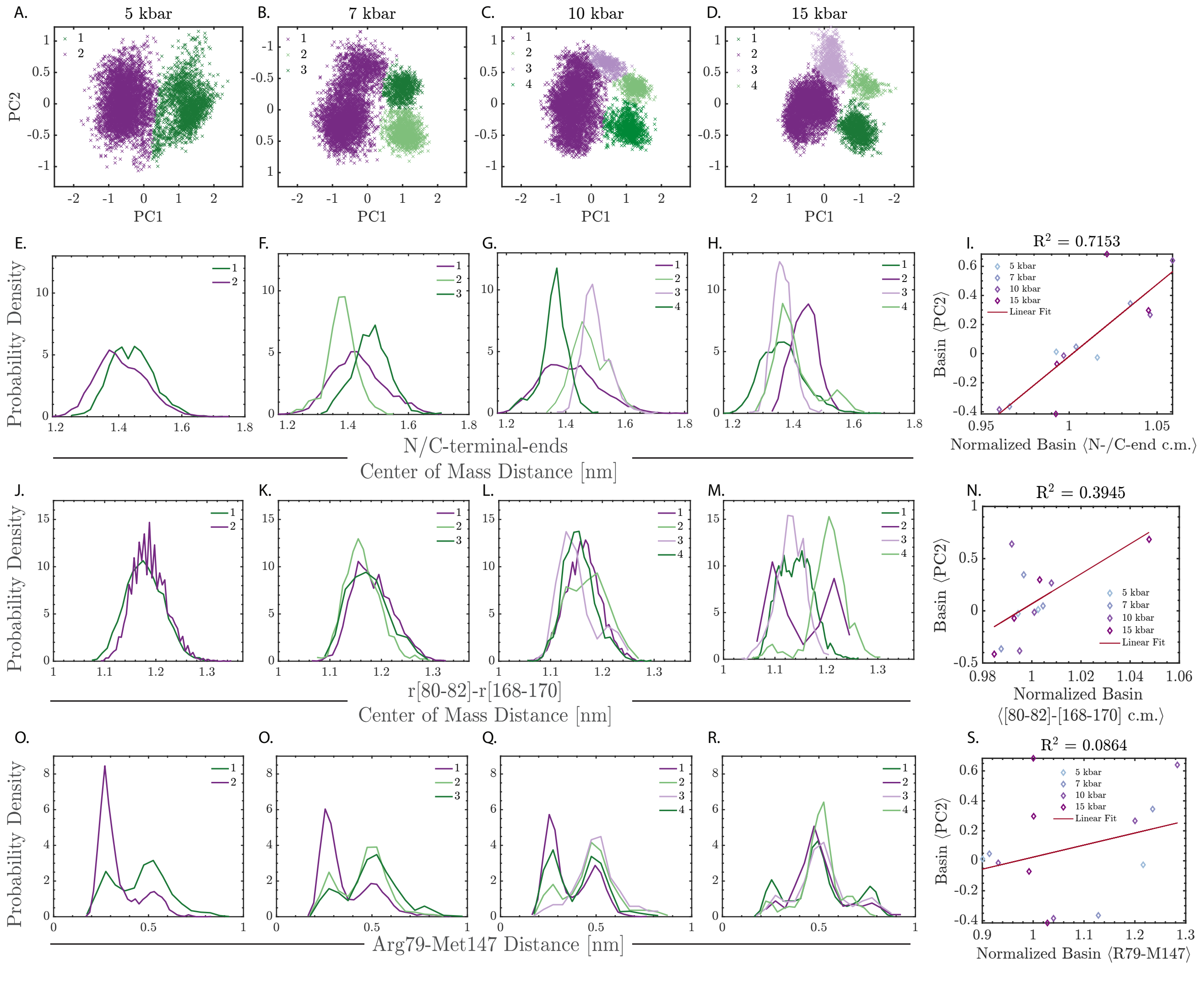


FIG. S3. (A-D) Distinct free energy basins within the PC1-PC2 space colored based on their association with different principal component projections: green and purple for PC1 and dark and light for PC2. (E-H, J-M, O-R) Conditional histograms of (E-H) N-/C-td ends center of mass distance, (J-M) residue [80-82]-[168-170] center of mass distance, and (O-R) R79-M147 distance for the basins distinctly colored in (A-D). (I, N, S) Linear correlations between the normalized mean (I) N-/C-td ends center of mass distance, (N) residue [80-82]-[168-170] center of mass distance, and (S) R79-M147 distance with mean PC2, respectively.

### S3. HYDROGEN BONDING

In order to probe the effect of high pressure on the solvation of the hydrophobic core, we enumerate the number of hydrogen bonds formed by the six hydrophobic residues of the interdomain interface. We adopt a geometric definition of hydrogen bonding, wherein two entities are identified as hydrogen-bonded if: (i) the donor-acceptor distance is smaller than 0.35 nm, (ii) the donor-acceptor-hydrogen angle is smaller than 30° [1]. We consider OH and NH groups as donors and nitrogen and oxygen atoms as acceptors, and distinguish hydrogen bonds formed between two intramolecular functional groups (within the protein) and those formed between a functional group and water molecules. Figure S4 depicts the pressure dependence of the histograms of the number of intramolecular (Fig. S4A) and solvent-protein (Fig. S4A) hydrogen bonds. While the intramolecular hydrogen bond distribution is virtually insensitive to pressure, there is a slight rightward shift of the protein-solvent hydrogen bond distribution. More precisely, the mode of the latter shifts from 5 (at ambient pressure) to 6 (at high pressures) consistent with the observed intrusion of the hydrophobic core by water molecules (depicted in Fig. 7C).

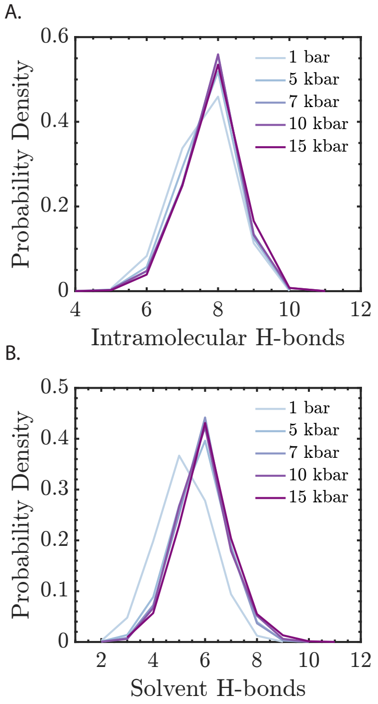


FIG. S4. Histograms of the number of (A) intramolecular (protein-protein) and (B) protein-solvent hydrogens bonds formed by the six hydrophobic residues of the interdomain interface as a function of pressure.

[1] D. Van Der Spoel, P. J. Van Maaren, P. Larsson, and N. Tîmneanu, Thermodynamics of hydrogen bonding in hydrophilic and hydrophobic media, J. Phys. Chem. B **110**, 4393 (2006).
